# Heterogeneity induced block co-polymer segregation in confinement

**DOI:** 10.1101/2024.10.31.621393

**Authors:** Dibyajyoti Mohanta, Manish Dwivedi, Debaprasad Giri

**Affiliations:** Department of Chemistry, University at Buffalo, 476 Natural Sciences Complex, Buffalo, New York 14260, USA; Department of Physics, Institute of Science, Banaras Hindu University, Varanasi 221005, India; Department of Physics, IIT (BHU), Varanasi 221005, India

## Abstract

Motivated by the work on block copolymer models that provide insights into epigenetics driven chromosome organization, we investigate the segregation behavior of five distinct 2-block co-polymers (BCPs) system with varying block sizes, confined within both symmetric and lateral geometries. Using exact enumeration method and Langevin dynamics simulation, our simple self-avoiding polymer model reveals robust behaviors (across statics and dynamic studies) despite strong finite-size effects. We observe that as block length increases, polymer compaction intensifies relying on non-specific interaction, leading to longer segregation times. The dynamic study clearly demonstrates the formation of globular lamellar phases and condensed, stable complex structures in long-range block copolymer (BCP) systems, providing a simplified analogy to lamellar-mediated chromatin compaction, which involves structures that are difficult to segregate under physiological conditions. Dominance of specific interaction over non-specific interaction in long range BCP systems leads to phase separation driven self assemblies which provides a simplified analogy to heterochromatin—inactive or stable domains. In contrast, short-range block sequences remain in a coiled state, exhibiting minimal overlap or interaction due to strong short range attraction, which may corresponds to euchromatin regions where diverse epigenetic states coexist, resulting in active, non-condensed structures. We also observe that asymmetric or lateral confinement favors more segregation between the BCPs irrespective of their underlying sequence.

## I. INTRODUCTION

The complete genome sequence, extending to roughly the length of a couple of meters, is packaged into the nucleus, a small dedicated compartment in living cell (size of few *μ*m) [1, 2]. At first thought, one might assume that the extensive folding and packaging of DNA would lead to an impenetrable and disordered configuration. Over the decade, studies have demonstrated that genomes exhibit intricate organization through fractal and hierarchical patterns, although the precise mechanisms driving this structure remain unclear[3, 4]. One of the key mechanisms leading to this specific (cell-dependent) genome folding is epigenetics [5, 6]. Epigenetic marks do not alter the underlying DNA sequence, and can be thought of as an additional level of information beyond the DNA sequence[5].

In the past decade, the problem of genome folding and the role of epigenetics has also attracted the attention of theoretical physicists [7–11]. Interestingly, the problem of genome folding shares natural similarities with the organization of block co-polymers (BCPs) under confinement [12]. Although challenging, recent coarse-grained polymer models for genome organization have sought to relate the distribution of epigenetic marks to block co-polymers by depicting them as block co-polymers with ‘colored beads’(Fig.1) [13, 14]. A so-called “block co-polymer” (BCP) is defined as a self-avoiding walk polymer consists of two or more kinds of monomer with periodic/random blocks (successive monomers of same kind), with a nearest-neighbour interaction between the specific monomers (say, A-A or B-B) or non-specific monomers (say A-B). The specific *ϵ*_*AA/BB*_ (non-specific *ϵ*_*AB*_) interactions thus play role in interaction driven polymer organization/ folding.

**FIG. 1:**
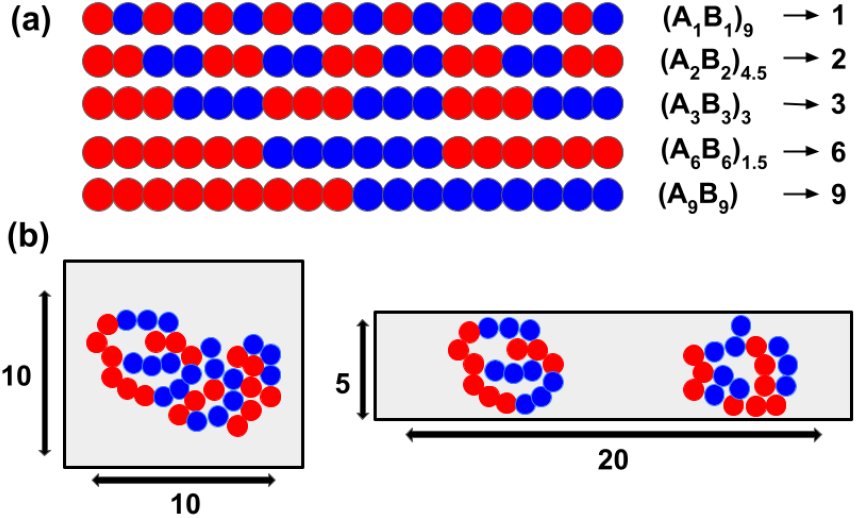
(a) shows the schematic presentations of five different block co-polymers (*A*_1_*B*_1_)_9_, (*A*_2_*B*_2_)_4.5_, (*A*_3_*B*_3_)_3_, (*A*_6_*B*_6_)_1.5_, *A*_9_*B*_9_ which are presented in the text as 1, 2, 3, 6 and 9 respectively in the 2-polymer system. Polymer 2 in each different systems are complementary sequence to the polymer 1 and given by (*B*_1_*A*_1_)_9_, (*B*_2_*A*_2_)_4.5_, (*B*_3_*A*_3_)_3_, (*B*_6_*A*_6_)_1.5_, and *B*_9_*A*_9_ respectively (not shown in the figure). (b) represents the schematic representations of 2 block co-polymers (3-3’ system as an example) within square confinement *S*_*A*_ with side length *L*_*xA*_ = *L*_*yA*_ = 10 units and rectangular or lateral confinement *S*_*B*_ with dimensions of length *L*_*xB*_ = 20 and height *L*_*yB*_ = 5.

Multiple studies using simple yet varied block co-polymer models have provided valuable insights into chromatin folding, TAD loop formation, and SMC protein complex-mediated domain segregation. Daniel Jost et al. used a minimal block co-polymer model, ((*A*_10_*B*_10_)_6_), to show that specific interactions favor small domain formation, while non-specific interactions allow global compaction [15]. Luca Fiorillo et al. demonstrated that block co-polymer models successfully describe A (or B) compartmentalization and Topological Associated Domains (TAD) formation [16]. Simona Bianco et al. [17] employed block co-polymer and strings- and-binders formalism to reflect distant contact formation in DNA sequences (looping). Additionally, Michieletto et al. [7] used a dynamic Potts model on a homopolymer, transforming it into a recolorable heteropolymer or block co-polymer model. They showed that this model can describe the transition from a disordered swollen state at higher temperatures to a compact ordered state at lower temperatures. Most of these studies focus on isolated polymer models, emphasizing self-interaction-induced polymer folding to provide insights into chromatin folding. However, the dynamics of multiple block copolymer models in polymer organization remain underexplored, especially in finite-size systems where the presence of one block copolymer can significantly influence global and local structural changes. Also, entropy, as a measure of randomness, can paradoxically drive the ordered organization of biopolymers when subject to strong asymmetric confinement [18]. Biologically relevant confinements (such as chromosome domains in different phases) vary shapes and dynamically change volume fractions, which can significantly influence the structural transitions of polymers, including their collapse, folding, or unfolding [19, 20]. One of the aim of this article is to investigate how strong confinement with varying shape affects the organization of block copolymers, particularly in the context of phase behavior and structural patterns.

Enthalpic contributions arising from specific and non-specific interactions also compete with entropy to drive the spatial organization of polymers under confinement [21]. Specifically interacting segments tend to compartmentalize into compact, energetically favorable domains, often occupying peripheral regions of the confined space, while non-specifically interacting monomers are typically found in the non-dense central region of the nucleus [15, 22]. The stiffness and rigidity of these polymers are strongly modulated by the nature of their interactions. Co-polymers with short-range repeat sequences are more likely to form rigid, extended structures due to localized enthalpic interactions, whereas those with long-range repeats favor the formation of flexible smaller domains within each repeating unit [23]. These short-range interactions can facilitate linear overlapping, which are hypothesized to play a critical role in processes such as chromosome de-condensation thereby enabling dynamic cellular functions. On the other hand, long block co-polymers can interact over many blocks leading to global compaction[24, 25]. Motivated by above, this model study aims to detail the organization of BCPs by varying repeat sequence lengths and interaction strengths while the monomer kind ratio remains same. We use a simple minimal model of two block co-polymers with varying sequences (*A*_1_*B*_1_, *A*_2_*B*_2_, *A*_3_*B*_3_, *A*_6_*B*_6_, and *A*_9_*B*_9_) to investigate the emergence of short domains, or global loop formation by varying specific and non-specific interactions. The core idea behind this study lies on the fact that the entropic and enthalpic contribution of confined block co-polymer compete with each other where entropic loss due to confinement always results in an increase in free energy, but coupled with energetic contribution due to specific or non-specific interaction or solvent property can increase or decrease the free energy depending on the nature of interaction. We used exact enumeration method and Langevin molecular dynamics simulation to study the statics and dynamics of the finite size block co-polymers in strong confinement.

The aim of the current study is two-fold. First, we examine the polymer organization through contact distribution using both static and dynamic analyses. In the second part, we investigate the size distribution of block copolymers and compare the results from static and dynamic studies to demonstrate and discuss the robustness of the findings. We also explore the free energy landscape of the two-polymer system and analyze the segregation time using the Fokker-Planck formalism. Lastly, we summarize our findings and discuss some limitations of the static analysis in the conclusion section.

## II. MODEL AND METHOD

We model two polymers, each having *N* + 1 = 18 monomers (chain length *N* = 17), as *self(mutual)-attracting-self(mutual)-avoiding walks (S(M)AS(M)AW)* within a square box confinement *S*_*A*_ with each side of length *L*_*xA*_ = *L*_*yA*_ = 10 lattice units, and a rectangular confinement *S*_*B*_ with dimensions of length *L*_*xB*_ = 20 and height *L*_*yB*_ = 5, both having an area of 100 square units maintaining same volume fraction 0.36 (Fig 1). The studied volume fraction aligns with the range of chromatin densities reported in the literature (e.g., Drosophila studies)[26–29]. Building on the ideas from [18, 21], we aim to investigate how the shape of the confinement (square versus rectangular) influences the organization of two block copolymers (BCPs), while keeping the total confinement area constant (see supplementary text for detailed simulation procedures).

We have considered five different block co-polymer systems with varying block sizes, where the block represents the repeating unit of monomers along the chain. All the BCPs in our model (see Fig 1 (a)) are composed of two types of monomers, A and B. System 1-1’ has block size *A*_1_*B*_1_ in polymer 1 (1) and *B*_1_*A*_1_ in polymer 2 (1’), with the sequences represented as (*A*_1_*B*_1_)_*n*1_ and (*B*_1_*A*_1_)_*n*1_, where *n*1 = 9 indicates the number of blocks/repeating units (block size of 2 monomers) in the 1-1’ system. Similarly, systems 2-2’, 3-3’, 6-6’, and 9-9’ consist of polymers (*A*_2_*B*_2_)_*n*2_ and (*B*_2_*A*_2_)_*n*2_, (*A*_3_*B*_3_)_*n*3_ and (*B*_3_*A*_3_)_*n*3_, (*A*_6_*B*_6_)_*n*6_ and (*B*_6_*A*_6_)_*n*6_, and (*A*_9_*B*_9_)_*n*9_ and (*B*_9_*A*_9_)_*n*9_, where *n*2, *n*3, *n*6, and *n*9 are 4.5, 3, 1.5, and 1 respectively. For all systems, the overall A and B monomer ratio is kept at 1, although the A and B monomer ratio in each polymer may vary (e.g., for the 6-6’ system, 6 has a ratio of 2/1, and 6’ has a ratio of 1/2).

The thermodynamic properties of such systems containing two BCPs chains are expressed in terms of averages of the physical observables derived from the partition function. The canonical partition function for the polymer chains of length *N* is expressed as:

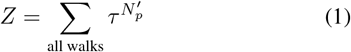

The summation is over all possible walks, and *τ* = exp(−*βϵ*) is the Boltzmann weight corresponding to any non-bonded nearest neighbor pair. *β* = 1*/*(*k*_*B*_*T*), where *k*_*B*_ is the Boltzmann constant and *T* is the temperature. 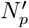 is the total number of nearest neighbor pairs, where each pair has energy *ϵ*.

We study the above system using a canonical ensemble approach where the partition function of the two-polymer system can be expressed as:

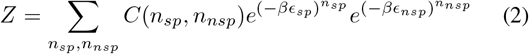

where *C*(*n*_*sp*_, *n*_*nsp*_) represents all possible conformations of two polymers with a total of *n*_*sp*_ specific contacts formed between any two monomers of same kind such as A-A or B-B, and *n*_*nsp*_ non-specific contacts formed between different monomers A and B. Here, both *n*_*sp*_ and *n*_*nsp*_ involve intra and inter contacts between the polymers with interaction energies *ϵ*_*sp*_ and *ϵ*_*nsp*_ respectively. In this study, we use reduced units with *k*_*B*_ set to 1 and temperature *T* fixed at 0.5. We vary the two interaction energies *ϵ*_*sp*_ and *ϵ*_*nsp*_ to study the five different BCP systems within confinements *S*_*A*_ and *S*_*B*_.

The exact partition function of this system can be calculated using the exact enumeration method, where all possible conformations are generated for the two block co-polymers of chain length *N*. However, as *N* increases, the number of conformations *C*(*n*_*sp*_, *n*_*nsp*_) increases rapidly within confinement for the studied volume fraction range (≈ 0.36) [30, 31].

The free energy of the two-polymer system within the confinement *S*_*A*_ or *S*_*B*_ can be expressed as:

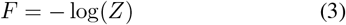

We have carried out langevin dynamics simulation by modelling our block co-polymer with beads that are interconnected with springs. The bonded monomers in the chain interact *via* finite extensible nonlinear elastic (FENE) potential and excluded volume interaction are modeled by repulsive part of Lenard-Jones (LJ) potential given by the following equations respectively:

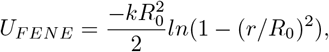

where *R*_0_ is the maximum extension between two consecutive beads, *k* is spring constant.

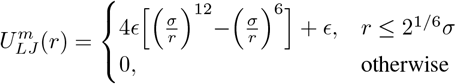

where, *ϵ* is the strength of the potential. The cutoff distance is set to be *r*_*min*_ = 2^1*/*6^*σ*.

The specific interaction (*AA*) and (*BB*) and non specific interaction (*AB*) between monomers is modelled by the LJ potential with a cutoff of 2.5*σ* and interaction energy *ϵ*_*AA*_, *ϵ*_*AA*_ and *ϵ*_*AB*_.

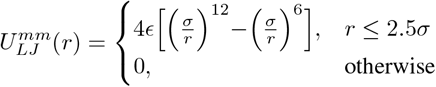

where, *ϵ* is the strength of the potential. The interaction between the walls of confinement and the polymer’s monomers 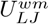 is modelled by the same repulsive part of LJ potential as the excluded volume interaction so 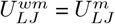.

The dynamics of the system is carried out using the following Langevin equation of motion:

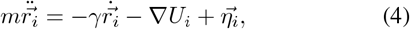

here *m* represents the mass of a monomer, *U*_*i*_ denotes total contribution from (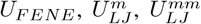 and 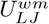) potential experienced by the monomer, and *γ* is the friction coefficient. The random force *η*_*i*_ follows a white noise characteristics *i*.*e. < η*_*i*_(*t*)*η*_*j*_(*t*^*′*^) *>* = 2*k*_*B*_*Tγδ*_*ij*_*δ*(*t − t*^*′*^), where *k*_*B*_ is Boltzmann’s constant and *T* is the temperature. The equation of motion is solved using a fifth-order predictor corrector method. Throughout the simulations, we work in reduced units, where *ϵ*_*AA*_ = 1.0, *ϵ*_*BB*_ = 1.0, *σ* = 1.0 and *m* = 1.0, are the units of energy, distance, and the mass of the beads, respectively. Other quantities are measured in units of 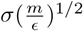. We used parameters *R*_0_ = 1.5*σ, k* = 30, *k*_*B*_*T* = 0.5 and *γ* = 0.7. The time step ∆*t* is taken to be 0.001[32].

## III. RESULTS

### A. Statics: Mutual Contact distribution

We quantify the degree of compaction or mixing between two polymers using a single parameter in the static study: the number of mutual contacts (*n*_*c*_) where a mutual contact is a inter-chain contact [21]. When the number of mutual contacts reaches its maximum value 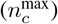, which may exceed the total number of monomers *N* = 18 in a polymer, the polymers are considered fully mixed. Conversely, when *n*_*c*_ = 0, the polymers are completely segregated, with no mutual contacts between them. We aim to investigate heat maps of mutual contacts as a function of specific (*ϵ*_*sp*_) and non-specific (*ϵ*_*nsp*_) interaction energies. This will also help us to understand which interaction energy (specific or non-specific) can play decisive role in segregation in terms of zero mutual contacts.

Figure 2 shows the heat plots of mutual contacts (*n*_*c*_) between the two polymers for different systems, with the upper panel representing the case within *S*_*A*_ and the lower panel for *S*_*B*_. At near *ϵ*_*sp*_ ≈ 0 and *ϵ*_*nsp*_ ≈ 0, all different block co-polymer systems exhibit similar behaviour, as they lack attractive strength between constituent monomers and behaves as non-interacting self avoiding walks (SAWs) with zero mutual and individual contacts (see Appendix and Fig. 6). The polymers near this interaction range, due to only entropic contributions, discourage any overlapping (which entails an entropic cost). Consequently, they are mutually independent (non-overlapping) solely relying on configurational entropy. Bertrand R. Care, et. al. [33] reported similar trend where block co-polymer remain in coil form when both specific and non-specific attractions remain very low.

**FIG. 2:**
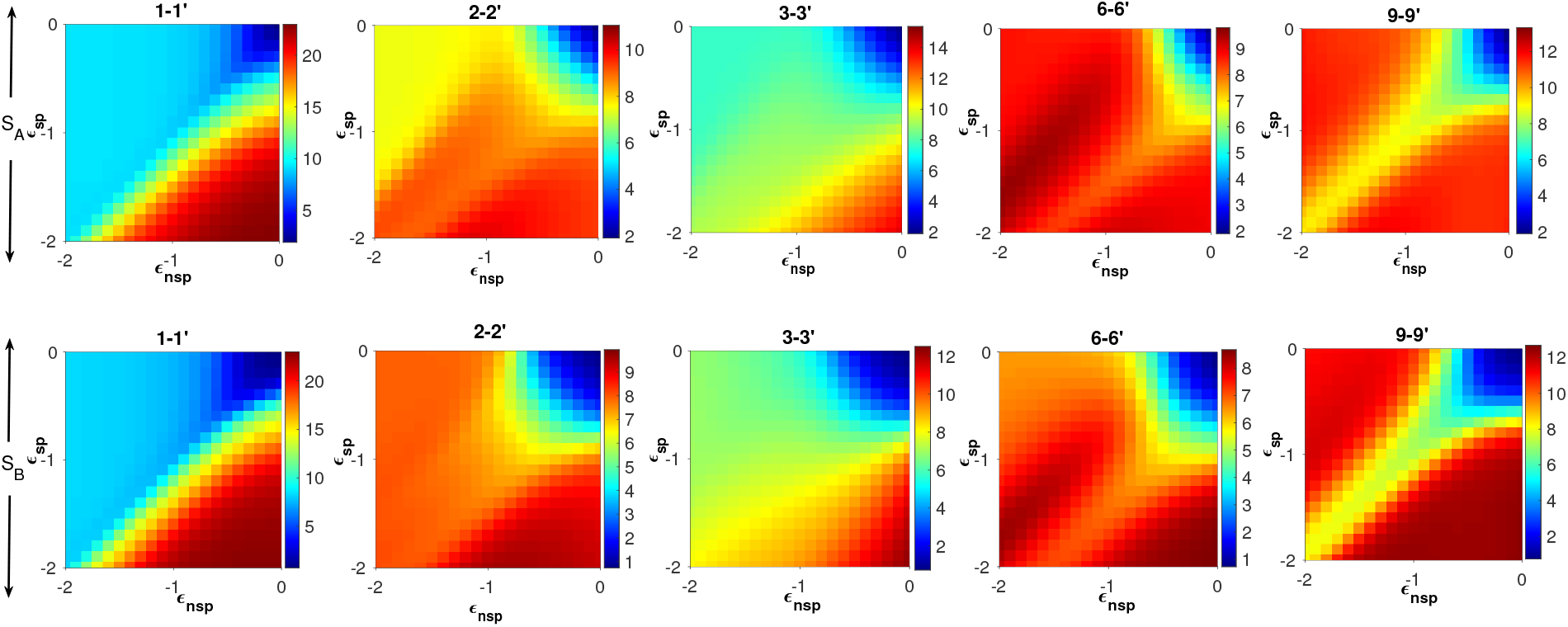
The horizontal upper panel shows mutual contacts heatmap for shape *S*_*A*_ and lower pannel presents the mutual contacts heatmap as a function *ϵ*_*AA*_ and *ϵ*_*AB*_ for shape *S*_*B*_

1-1’ system in both *S*_*A*_ and *S*_*B*_ shows that the mutual contacts remain low within the range − 2 *< ϵ*_*nsp*_ *<* 0 and − 0.5 *< ϵ*_*sp*_ *<* 0, indicating that at these interaction energies, neither is strong enough to overcome the entropic freedom of polymers, resulting configurations with less overlap. D. Jost et.al. [15] also observed similar behavior where block co-polymers remain in coil form or extended form when both specific and non-specific interaction strength remains low. However, at lower *ϵ*_*sp*_ *<* − 1.0, even when *ϵ*_*nsp*_ = 0, there is a clear evidence of polymer overlap, as mutual contacts increase to ≈ 22. This was earlier reported in lattice studies where short range sequence polymers form parallel coil configurations overlapped with each other [34]. Now, if instead *ϵ*_*nsp*_ is decreased, the polymers initially remains overlapped, but as *ϵ*_*nsp*_ *< ϵ*_*sp*_, mutual contacts decrease sharply. This suggests that 1 and 1’ prefer to overlap when *ϵ*_*sp*_ dominates over *ϵ*_*nsp*_. This behavior can be regarded as a limitation of lattice study as due to the alternating pattern of 1 and 1’ in the square lattice, no *n*_*sp*_ intra-contacts are formed within polymers 1 and 1’ when *ϵ*_*sp*_ dominates(see Appendix and Fig 6, where intra contacts are zero near *ϵ*_*nsp*_ = 0 for 1-1’). The contribution to *n*_*sp*_ comes solely from only mutual contacts *n*_*c*_, such that *n*_*sp*_ ≈ *n*_*c*_. Therefore, when *ϵ*_*sp*_ *< ϵ*_*nsp*_, polymers 1 and 1’ come closer, forming more overlapping contacts (inter contacts), minimizing the system’s free energy. When *ϵ*_*nsp*_ *< ϵ*_*sp*_, the polymers 1 and 1’ favor forming *n*_*nsp*_ contacts, where both intra- and inter-polymer non-specific interactions contribute to the total energy. Since it is entropically unfavorable for them to remain overlapped when both intra- and inter-contacts formation are available, polymers 1 and 1’ both form individual coils causing mutual contacts *n*_*c*_ to decrease sharply (see Appendix, 1-1’ case). The non-specific interaction driven self association can be seen as a phase separated domain of A-B loops or can be termed as polymer-polymer phase separation and previously seen in short sequence studies [35]. Thus, stronger non-specific interaction strength *ϵ*_*nsp*_ discourages overlapping or compaction for 1-1’. One thing to note here is that, in the heatmap for the ‘1-1’ system of alternating A-B monomers, the parameter sets (*ϵ*_*nsp*_ = −1.0, *ϵ*_*sp*_ = 0) and (*ϵ*_*nsp*_ = 0, *ϵ*_*sp*_ = − 1.0) should feature the same behavior. However, as described, due to lattice limitations, we observe higher mutual contacts for (*ϵ*_*nsp*_ = 0, *ϵ*_*sp*_ = − 1.0) (see Fig. 6). We have not seen any qualitative difference in mutual contacts heatmap for *S*_*B*_.

The maximum overlap scenario differs substantially for 2-2’ relative to 1-1’, with the maximum *n*_*c*_ decreased to approximately 9 −10 at *ϵ*_*nsp*_ ≈ 0 and *ϵ*_*sp*_ ≈ − 2, while for the rest of the energy spectrum, the overlap remains relatively stable in *S*_*B*_. This suggests that polymers 2 and 2’ tend to remain in individual coils, which maintain a limited number of sharing contacts (crosstalk) throughout the energy range (see Appendix 2-2’ heat maps). The higher individual contacts (appendix) within *S*_*A*_ for 2-2’ system at *ϵ*_*nsp*_ ≈ −2 and *ϵ*_*sp*_ ≈0 restates that confinement *S*_*B*_ disallows nesting of globules and introduces more entropic stabilization in *S*_*B*_ by linear overlapping [36]. In other words, the non-varying 2-2’ heatmaps indicates that the 2-2’ system might be in a phase-locked state or metastable configuration in both confinement scenarios especially in *S*_*B*_. There has been a numerous studies where researchers have found meta stable sates or multiple minimas throughout the configuration space at intermediate energy interactions [15, 37, 38]. For the 2-2’ polymer system, confinement introduces noticeable differences between *S*_*A*_ (upper panel) and *S*_*B*_ (lower panel), as there is significantly less variation in ⟨*n*_*c*_ ⟩ across the energy spectrum in *S*_*B*_ compared to *S*_*A*_.

In the 3-3’ polymer system, the region with high overlap (*n*_*c*_ ≈ 12 −13) at *ϵ*_*sp*_ *< ϵ*_*nsp*_ shrinks. This indicates that polymers 3 and 3’ generally prefer less overlap, maintaining mutual contacts around *n*_*c*_ ≈ 8 throughout most of the energy phase diagram. Similar to the 2-2’ system, there is no sharp transition in *n*_*c*_, and the cross talk between 3 and 3’ remains with a finite value throughout the energy spectrum. Notably, in the 3-3’ system, mutual overlap near zero non-specific interaction energy is primarily driven by specific interactions. Additionally, at the highest degree of mixing, both confinement scenarios show nearly the same range of mutual contacts with both polymers forming individual globules, suggesting that the overlap is not purely linear (see Appendix and FIg. 6).

The 6-6’ case is the most asymmetric, where the A/B ratio across the entire two-polymer system is 1, but differs within each individual polymer (A:B ratios are 2:1 and 1:2 for polymers 6 and 6’, respectively). Across a range of *ϵ*_*nsp*_ values, the number of mutual and sum of individual contacts (6 + 6^*′*^) remain relatively stable at approximately 9 and 16-18 respectively for each specific *ϵ*_*sp*_ (see 6 6-6’ map). The mutual contact map for the 6-6’ system suggests a stable formation with loose cross talk between 6 and 6’, making it one of the stable overlapped system among the cases studied. A slight non-linearity in *n*_*c*_ is observed as *ϵ*_*nsp*_ decreases, indicating potential structural rearrangements or emergence of minimas [15, 37]. For example, polymers 6 and 6’ may overlap along their specific contacts when *ϵ*_*nsp*_ ≈ 0, while at higher non-specific interaction strengths, they may adopt individual 𝒮-shaped structures that still share few non-specific contacts to maximize *n*_*nsp*_. Emergence of complex shapes such as 𝒮, 𝒱, 𝒰, etc. in block co-polymer studies were also seen in previous studies of Das et. al, Stefano, et.al [39, 40]

In the final case of polymers 9 and 9’, the system features significant mixing across the studied interaction energy range. Similar to the 6-6’ system, it shows stable contact distribution but with a higher degree of overlapping. At very high specific interaction strengths, *S*_*B*_ shows more overlapping and less individual folding (folding into itself), suggesting that the polymers may rely on specific contacts for overlap (see 6 in Appendix). This is well discussed in previous findings of polymer models led understanding of chromatin organization where block co-polymers are micro phase separated at higher specific interactions [15, 40, 41]. Furthermore, in the homopolymer regime where *ϵ*_*sp*_ = *ϵ*_*nsp*_, mutual over-lapping decreases significantly as the polymers behave more like entropic objects, leading to a loss of mutual contacts and increase in individual folding (Appendix) [21]. Unlike the other systems, decreasing *ϵ*_*nsp*_ does not significantly impact the mixing at higher non-specific interaction strengths. As non-specific interaction strength increases, both confinement scenarios show similar levels of mutual overlapping in terms of contacts, which suggests that the polymers may not be in a linear overlapped configuration (if linear overlap were the case, *S*_*B*_ would exhibit more mutual overlapping due to its lateral shape) unlike in higher specific strength regime. It indicates that the non-specific interaction strength leads both 9 and 9’ to form globule structures which was seen in the work of D. Jost et. al at higher heterogeneity interaction strength [15]. Considering the interplay between mutual and individual contacts, and the overall behavior of overlapping and segregation, we observe that as the length of the blocks increases the polymers become more miscible or overlapped with each other (except 1-1’ system where due to lattice restriction mutual overlapping are very high at low non-specific interaction strength). For the rest of this study, we fix *ϵ*_*sp*_ = − 1 while varying the non-specific interaction energy *ϵ*_*nsp*_ to explore the detailed quasi-equilibrium statics and dynamics of segregation as the system’s heterogeneity changes.

### B. Dynamics: Contact maps variation

The dynamic study validates our claims from the static analysis while extracting additional insights on the evolution of 2-polymer systems under varying non-specific interaction energies, with the specific interaction energy fixed at *ϵ*_*sp*_ = 1. To this end, we present the contact probability maps, depicting intra-(individual) and inter- (mutual) contacts for different 2-polymer systems within *S*_*A*_ and *S*_*B*_ from molecular dynamics simulations shown in Figure 3.

**FIG. 3:**
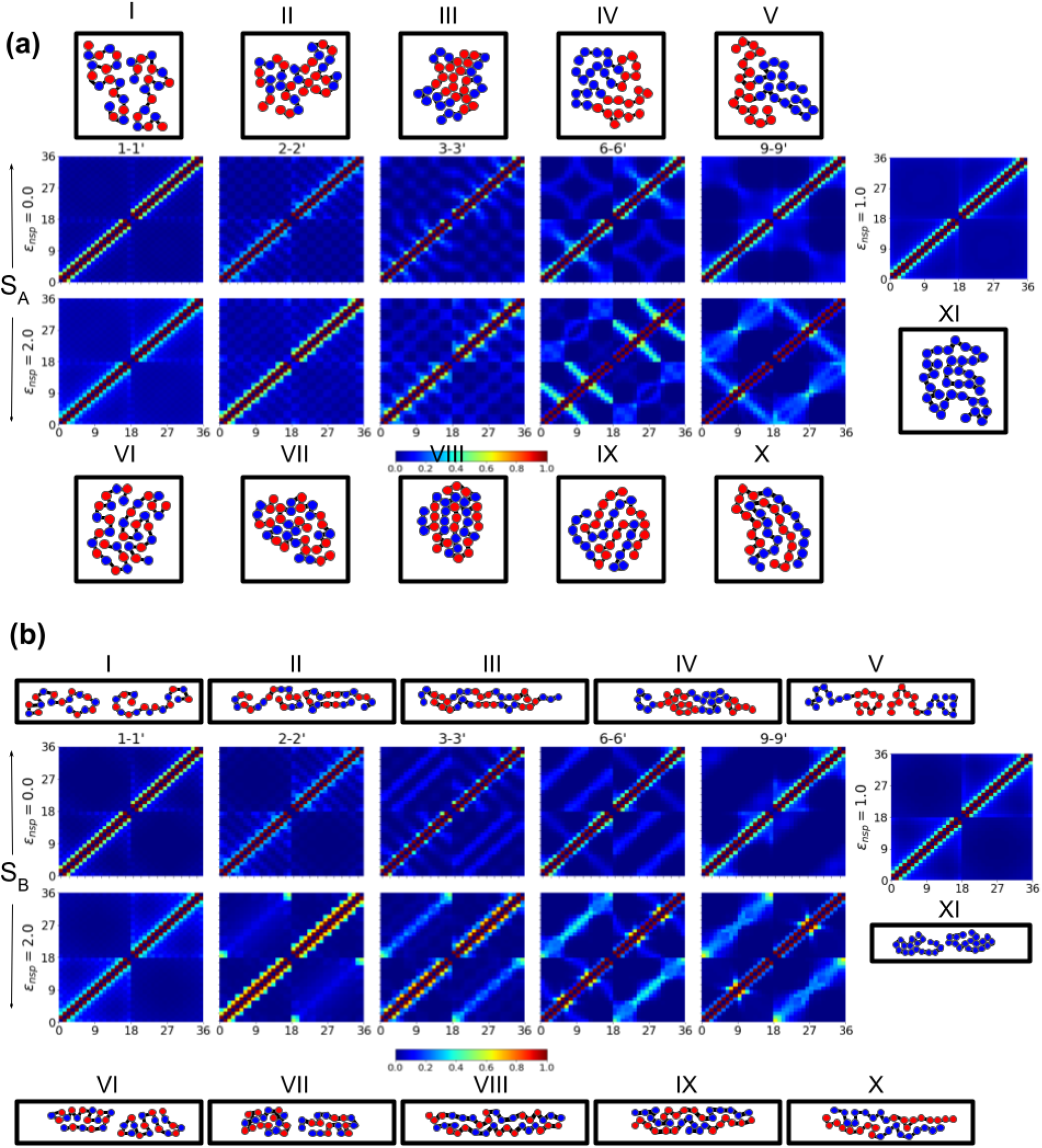
(a) shows the contact maps for different block co-polymer systems (1-1’, 2-2’, 3-3’, 6-6’, and 9-9’) within *S*_*A*_, where the x- and y-axes span from 1 to 36. Indices 1 to 18 correspond to polymer 1 (1, 2, 3, 6, and 9), and indices 19 to 36 correspond to polymer 2 (1’, 2’, 3’, 6’, and 9’). Each row of the maps represents a different non-specific interaction energy at *ϵ*_*nsp*_ = 0.0 and 2.0 (from top to bottom), with the specific energy fixed at *ϵ*_*sp*_ = 1.0. Each column corresponds to a different polymer pair (1-1’, 2-2’, 3-3’, 6-6’, and 9-9’, from left to right). The last column represents the case where *ϵ*_*nsp*_ becomes 1.0, making all systems identical, with both polymers behaving as homopolymers. For clarity, one of the dominant configurations is shown next to each contact heat map for the same set of energies. The contact maps are symmetric across the *x* = *y* line, reflecting that contacts between pairs (e.g., 1-1’ and 1’-1) are equivalent. (b) shows the contact maps for the same systems within *S*_*B*_.

For the 1-1’ studied case within the *S*_*A*_ geometry (Figure 3 (a)), we observe that both mutual overlap and individual folding probabilities remain consistently low for the upper heatmap *ϵ*_*nsp*_ = 0. This reveals that both 1 and 1’ prefers to be found in individual coils rather overlapping or forming globules due to the fact that the entropic cost would be higher of overlapping or folding while following the alternating specific inter or intra contact formation (at *ϵ*_*nsp*_ = 0 and *ϵ*_*sp*_ = 1.0) (see the next the configuration diagram Fig. 3 (a) I). At increased non-specific interaction *ϵ*_*nsp*_ = 2.0 (lower map), individual contact probability increases, evidenced by the subtle brightening near the *x* = *y* line, as well as at *x* = 18 and *y* = 18, indicating a potential for loop formation. Actually, the presence of short-range sequences in the 1-1’ system appears to hinder interactions, leading to frustration, with minimal folding or overlapping occurring throughout the dynamic regime. To examine the effect of confinement on segregation, we also plot the contact maps for *S*_*B*_ across the five different systems (see fig. 3 (b)). Under lateral confinement, the 1-1’ system at *ϵ*_*nsp*_ = 0.0 behaves similarly *S*_*A*_ but with more enhanced separation between them and a few intra contacts due to entropic stabilization provided by the confinement [36]. At higher interaction energy *ϵ*_*nsp*_ = 2.0, the polymers tend to fold more individually, resulting in a higher probability of contacts within each polymer and minimal mutual overlap between 1 and 1’ (see configuration Fig. 3 (b) VI). Notably, at *ϵ*_*nsp*_ = 0, where only the specific contact energy is dominant, individual contact probabilities are higher than mutual overlap for both *S*_*A*_ and *S*_*B*_. This suggests that, while in the static case 1 and 1’ cannot make individual contacts or fold relying upon specific interactions which should be considered a lattice artifact as claimed by the above section, the dynamic case allows for individual folding and segregation [15, 42]. So, unlike the static case where a clear mixing-demixing transition can be observed due to the lattice restriction, the dynamic study fully captures the competition between energy and entropy without any model limitation.

The 2-2’ system also exhibits low mutual contact probabilities, with minimal variation across different interaction energies. This behavior is consistent with the static analysis, suggesting that the 2-2’ system remains in a steady state with limited mutual contacts and finite individual folding. Throughout the energy range for *S*_*A*_, the checkerboard pattern shows formation of stickers and spacers of size 2 signifying micro-phase separation of like monomers which was earlier seen in block co-polymer studies [15]. The strong contact probability near the *x* = *y* line for *ϵ*_*nsp*_ = 2 arises due to the non-specific contact between the every closest B monomers to A monomers. This leads to kinks along 2 and 2’ which have resemblance to the pearl necklace feature of alternating block co-polymers [7, 43]. However, both specific interaction strength (*ϵ*_*nsp*_ = 0 and *ϵ*_*sp*_ = 1, Fig. 3(a) II) and non-specific interaction strength dominance (*ϵ*_*nsp*_ = 2 and *ϵ*_*sp*_ = 1, Fig. 3(a) VII) for the 2-2’ system lead to structures that rely more upon individual contacts, thereby leading to frustration or phase-locked conformations, as pointed out by the static study [44]. Many studies have observed and discussed frustration in heteropolymer models describing chromatin [15, 45]. For *S*_*B*_, the 2-2’ system shows no significant mutual overlap, and at *ϵ*_*nsp*_ = 2.0, the likelihood of small folds or kinks within the individual polymers 2 and 2’ gets stronger than *S*_*A*_ due to geometric restriction (see adjacent configuration of 2 and 2’). Additionally, there is a finite probability that 2 and 2’ make contacts primarily at their ends leading to long range loops, resulting in a higher contact probability at these locations, which suggests a linear configuration within *S*_*B*_ [46].

The 3-3’ system follows a similar trend to the 2-2’ system but with larger block sizes, leading to the emergence of a checkerboard pattern in the contact map. This pattern suggests that specific interaction energies at *ϵ*_*nsp*_ = 0 pre-dominantly govern the preferential contacts between blocks (AAA or BBB) leading to microphase separation of similar monomers. At *ϵ*_*nsp*_ = 2.0 (see Fig. 3(a) VIII)), individual folding becomes more pronounced, indicating an increase in non-specific contact formation as alternate blocks brighten in the heat-map. The checker board pattern reconfirms formation of folds by sticking of monomers of different types along the contour of the chain [42]. However, the blocks corresponding to mutual contacts remain less bright, reflecting the low probability of mutual overlap although there is a finite cross-talks between 3 and 3’. In the 3-3’ system under *S*_*B*_, at *ϵ*_*nsp*_ = 0, we observe a small but finite probability of shifted linear overlap between 3 and 3’ (shown in the adjacent configuration Fig. 3(b) III). As the non-specific interaction energy increases, the mutual overlap increases probability of having linear overlapped configuration increases. Additionally, there is a strong likelihood of small folds (thread like configuration with unknotted small folds) or kinks formation, driven by non-specific interactions in both confinements *S*_*A*_ and *S*_*B*_, as seen by the increased concentration in probability near the *x* = *y* line. The emergence of short as well as long folds suggests that both 3 and 3’ form linear, pearl necklace structures which were also found in earlier chromatin modelling studies [7, 43].

The 6-6’ and 9-9’ systems exhibit more intricate patterns compared to the earlier cases. At *ϵ*_*nsp*_ = 0.0, both individual folding and mutual overlap take place due to the formation of specific contacts leading to micro phase separation, though with relatively low probabilities [15, 47]. For both systems, we have seen specific interaction driven micro phase separation in both confinements (see Fig. 3(a) IV and Fig. 3(b) IV). The higher contact probability near *xy* = *c* (where c is constant) line within the individual contact map region suggests folds within same polymer and indicates that both polymers 6 and 6’ adopt 𝒮-shaped structures, with cross talks occurring along non-specific contact regions at *ϵ*_*nsp*_ = 2 (see adjacent configuration Fig. 3(a) IX). This configuration can be visualized as two 𝒮-shaped polymers in close proximity, sharing a few contacts with low probability, but not fully “hugging” each other (as evidenced by the lower intensity in the middle block compared to the outer overlapping blocks). These lamellar phases are result of varying interactions along the configurations space leading higher miscibility and has been emerged in similar studies over the years [40, 48**?** –50]. In the 6-6’ system (at *ϵ*_*nsp*_ = 2.0), lateral confinement in *S*_*B*_ restricts the formation of distinct 𝒮-shaped structures at higher non-specific interaction strengths. Instead, a finite degree of linear overlap between 6 and 6’ is observed in *S*_*B*_, a behavior less pronounced in *S*_*A*_ (Fig. 3(b) IX).

The 9-9’ system at *ϵ*_*nsp*_ = 0 within both *S*_*A*_ and *S*_*B*_, exhibits similar behavior, with both polymers 9 and 9’ forming internal folds due to specific contact formation and crosstalks between 9 and 9’ along the specific contacts region leads to micro phase separation (see Fig. 3(a) V and Fig. 3(b) V) [15, 50, 51]. This suggests that micro-phase separation leads to self-assembly and the formation of microdomains in long block copolymer systems. Such ‘local micro-phase separation’-induced aggregation has been previously observed in studies of polymer self-assembly forming micelles or in the aggregation of multiple smaller phase-separated clusters—a process analogous to how centromeres are organized in the nucleus [52, 53]. As *ϵ*_*nsp*_ increases to 2.0 within *S*_*A*_, the probability of individual contacts rises as well as the mutual overlap, as both polymers 9 and 9’ prefer to form concentratic 𝒰-shaped structures (“hugging”) the inner U aligned parallel to the outer one maintaining the non-specific contact formation leading to lamellar phase [48, 49]. Consistent with the static map of the 9-9’ system, we observe that the mutual overlapping probability is the highest among all the cases, and the overlapping pattern is of lamellar phase. H. Kasinsky, et. al. reported lamellar-mediated chromatin condensation in nuclei of algae [54]. The 9-9’ system in *S*_*B*_ prevents the formation of the specific 𝒰-shaped structure seen in other configurations. Instead, it favors a predictable linear overlap driven by non-specific contact formation [21]. These findings suggest that, despite the mixing observed in both the 6-6’ and 9-9’ systems, the lateral confinement of *S*_*B*_ inhibits the formation of specific shapes, leading to predominantly linear configurations due to confinement driven entropic stabilization [36]. These models, despite their simplicity, consistently demonstrate that the 9-9’ system is the most miscible while 1-1’ system is mostly segregated system, reinforcing the observation that larger block sizes promote increased miscibility.

Finally, we examine contact dynamics in the homopolymer regime, where both specific and non-specific interactions are set to 1 under confinements *S*_*A*_ (Fig. 3(a) XI) and *S*_*B*_ (Fig. 3(b) XI). In this regime, homopolymers behave as entropic objects dominated by intra-contact formation. We observe that polymers in *S*_*B*_ segregate more than those in *S*_*A*_, consistent with previous studies showing that lateral or asymmetric confinement promotes segregation [18, 21].

### C. Effect of sequence heterogeneity on the center of mass distribution of 2-polymer system

We examine and compare the separation between two polymers by analyzing the x-component of the center of mass (COM) differences in the confinements *S*_*A*_ (Fig. 4 a and b) and *S*_*B*_ (Fig. 4 c and d), using both static (Fig. 4 a and c) and dynamic (Fig. 4 b and d) studies. In the *S*_*A*_ system, the static analysis shows that the 1-1’ and 9-9’ pairs exhibit the least separation among the systems at *ϵ*_*nsp*_ = 0, whereas the dynamic study reveals that only the 9-9’ system, at low non-specific interaction strength making phase separation of similar kind monomers, leading to a low COM difference within *S*_*A*_ (Fig. 4 a and b) from both studies similar to the block copolymer studies of Jost et. al [15]. The 1-1’ static system behavior is explained in the heat map analysis where 1 and 1’ overlap strongly due to lattice limitation leading to decreased COM difference. The apparent discrepancy in the 1-1’ system is an artifact of the lattice was also observed by lattice studies of AB diblock polymer by A. Jayaraman et. al [34]. In the lateral confinement *S*_*B*_, the static study shows that all systems are in coil states with higher COM difference between them due to restricted configuration space. The dynamic study reinforces previous observations, showing that at low non-specific interaction strengths, most systems near *ϵ*_*nsp*_ = 0 form phase separated self assembled globules (Fig. 4 d), leading to higher ∆COMx (greater separation) [15]. So, in both confinements at low non-specific strength polymers remain in phase separated self organized loops or coils with greater separation in *S*_*B*_. When both interaction energies are equal (*ϵ*_*nsp*_ = *ϵ*_*sp*_), both static and dynamic studies show that all systems behave like homo polymers.

**FIG. 4:**
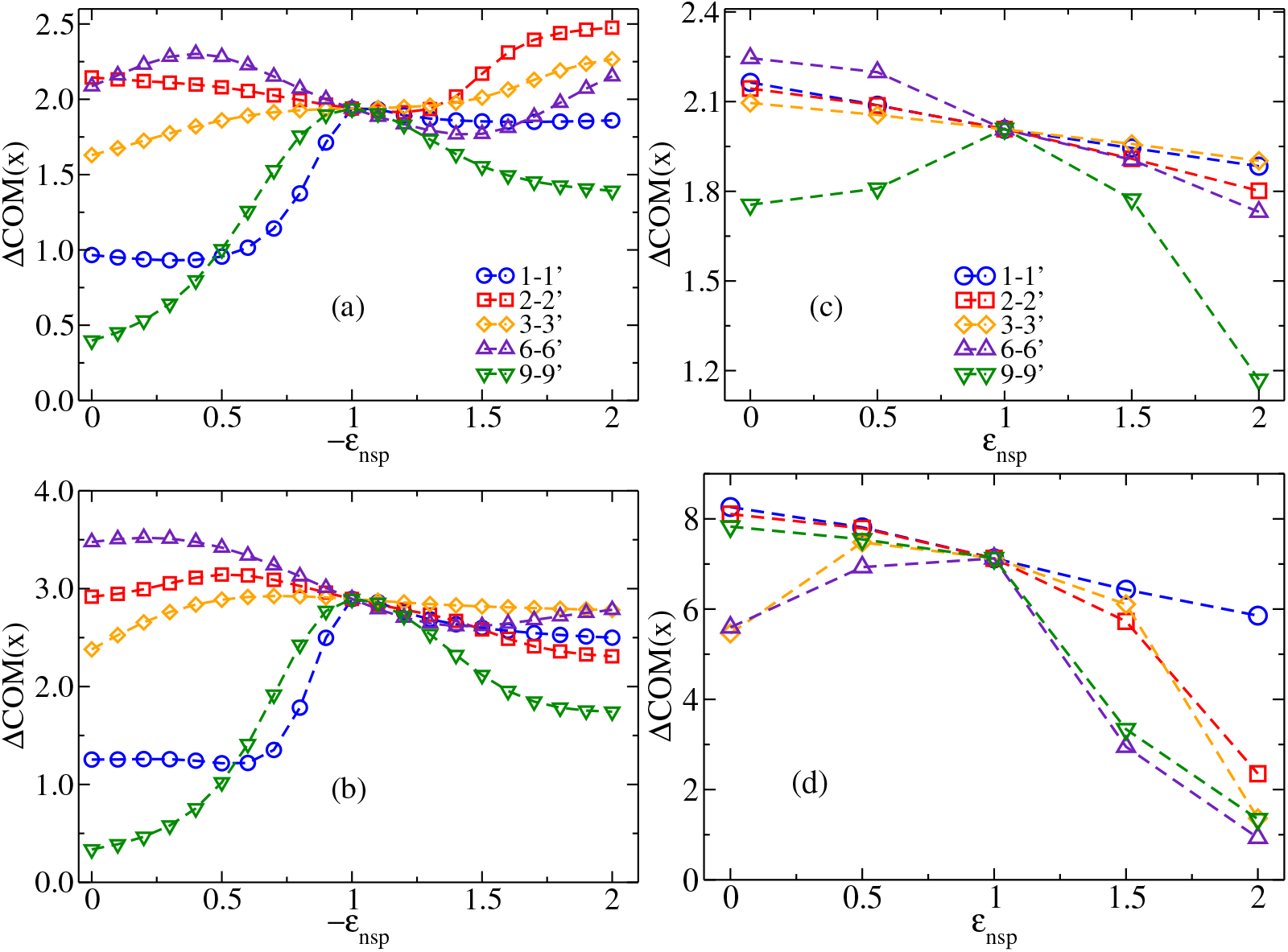
(a) and (c) represent the difference of x-component of Center of mass (∆COMx) between the two BCPs for the five different systems within *S*_*A*_. We compare ∆COMx from the static (a) and dynamic (c) studies. Similarly (b) and (d) represent ∆COMx within *S*_*B*_ where (b) and (d) are calculated from static and dynamic studies respectively

As the non-specific interaction strength increases within *S*_*A*_, the 9-9’ system becomes less separable from static study (Fig. 4 a) showing the robustness in result, as 9 and 9’ form “hugging” 𝒰-shaped structures. The other systems do not show significant variation in behavior and maintain a less average separation within *S*_*A*_. In the lateral confinement *S*_*B*_, at high non-specific interaction strengths, the static study shows gradual decrease in ∆COMx with similar trends of linear formation for all systems with lowest ∆COMx differences for 9-9’ system. The dynamic study clearly reveals that with increasing non-specific interaction strength the polymers are gradually coming close with long range BCPs such as 6-6’ and 9-9’ forming lamellar shapes (such as 𝒰 and 𝒮) [48, 49]. Additionally, we observed that at *ϵ*_*nsp*_ = 2.0, both 2 and 2^*′*^ form globules with shared contacts between them (this is also due to arrested micro phase separation) [55], while 3 and 3^*′*^ overlap linearly, resulting in reduced COMx difference along with the 6−6^*′*^ and 9−^*′*^ 9^*′*^ systems. It is important to note that, although the 2−2 system shows a smaller ∆COMx difference, it remains separated (see contact map of 2-2’ and 3 and 3^*′*^) as exhibit several kinks along the chain, which, despite the small ∆COMx, makes the 3 − 3^*′*^ system less compact or overlapped compared to the 6 − 6^*′*^ and 9 − 9^*′*^ systems. The quantitative difference in ∆COMx between *S*_*A*_ and *S*_*B*_ from both statics and dynamics study reveal that all polymers are more separated in lateral confinement *S*_*B*_ which is in accordance with the earlier studies [18, 21]. The ∆COMx analysis shows that at low non-specific interaction strengths, self-assembled micro-phase separation keeps long-range block copolymers close yet largely separated, whereas at high non-specific strengths, they mix together to form lamellar phases.

### D. Separation time

To investigate how polymer separation depends on specific sequences and whether there is a dependency on confinement, we calculate the segregation (separation) time using the Fokker-Planck formalism. This approach uses free energy as a function of mutual contacts *n*_*c*_ and can be expressed as

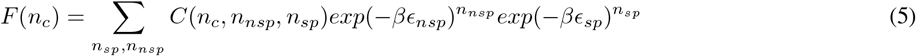

Using this information, at a specific *ϵ*_*nsp*_ with *ϵ*_*sp*_ fixed at −1, we calculate the partition function corresponding to all mutual overlaps (from 0 to 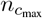) between the two polymers, where 0 contacts signify segregation and 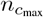represents maximum overlap in the static study. It is worth mentioning that the segregated scenario *n*_*c*_ = 0 is approximated by configurations that are separated only by the minimum COM difference with non mutual contacts between them. This ensures that configurations with larger COM differences (further apart) will not contribute to the time taken to segregate the polymers from *n*_*c*_ = 1 to *n*_*c*_ = 0, as they are already segregated. We also assume that segregation occurs very slowly, with mutual contacts detaching one at a time. At each stage, we calculate the free energy *F* (*n*_*c*_), which is then integrated to determine the segregation time using Fokker Planck formalism 6. Following the method developed in Ref. [56], we set *D*(*n*_*c*_) = 1. Since the present static study is confined to the lattice, the segregation time has been expressed in the following discrete form:

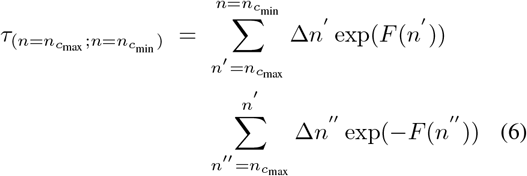

Here, the first summation adds the contributions of free energies from 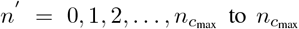, while the second summation sums up the free energy contributions from *n*^*′′*^ = 0 to *n*^*′*^. We have taken ∆*n*^*′′*^ = ∆*n*^*′*^ = 1 in lattice units.

The figure 5 shows that at zero non-specific interaction strength (*ϵ*_*nsp*_ = 0), the 1-1’ system exhibits the longest separation time in the static study (using the FP approach). The 1-1’ system in lattice remains highly overlapped which is earlier described as the lattice artifact, where polymers maximize *n*_*sp*_ contacts at *ϵ*_*nsp*_ = 0 and *ϵ*_*sp*_ = −1.0, relying only on overlapped contacts since individual folding cannot form any intra *n*_*sp*_ contacts in the lattice setup. Where as we can see that in dynamics there is a finite probability of forming loops through specific contact formation. Aside from 1-1’, the 9-9’ system also shows a longer separation time in comparison to other systems due to phase separation mediated moderate overlapping, as seen in the heat maps (Fig. 2 and Fig. 3) at *ϵ*_*nsp*_ = 0 for both confinements. In contrast, the 6-6’ system exhibits the least overlap (lowest *n*_*c*_) at *ϵ*_*nsp*_ = 0, due to pronounced individual folding into hairpin loops. This leads to the shortest separation or segregation time at *ϵ*_*nsp*_ = 0. In both confinement scenarios, the 2-2’ and 3-3’ systems show relatively short segregation times, though still longer than the 6-6’ system, as their short sequences favor individual folding (see Appendix and mutual heat maps).

**FIG. 5:**
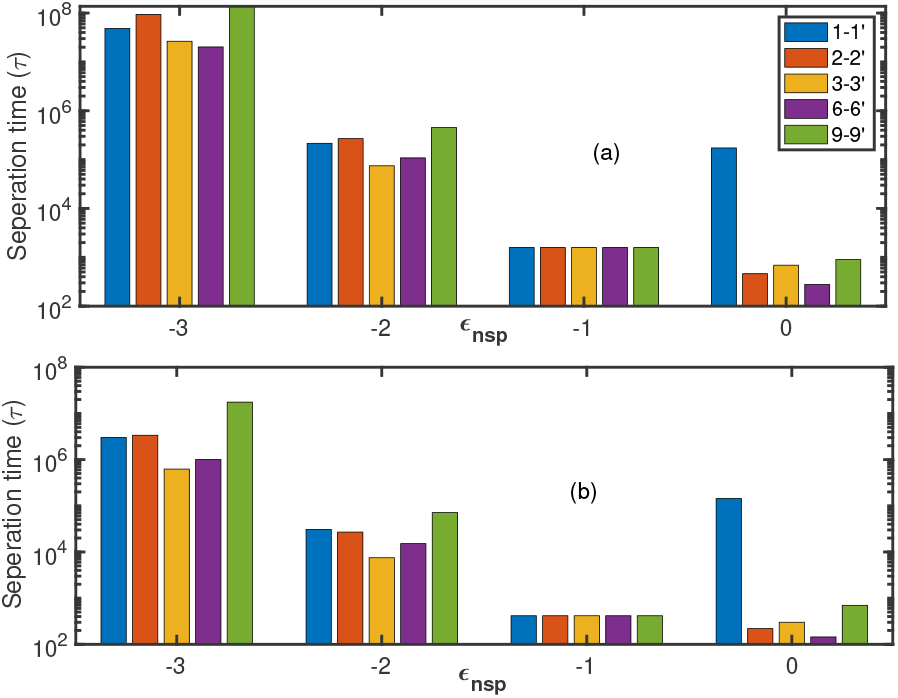
Fig (a) shows separation time calculated from FP formalism between poly 1 and poly 2 within *s*_*A*_ for five different BCP systems. (b) shows same within *S*_*B*_.

**FIG. 6:**
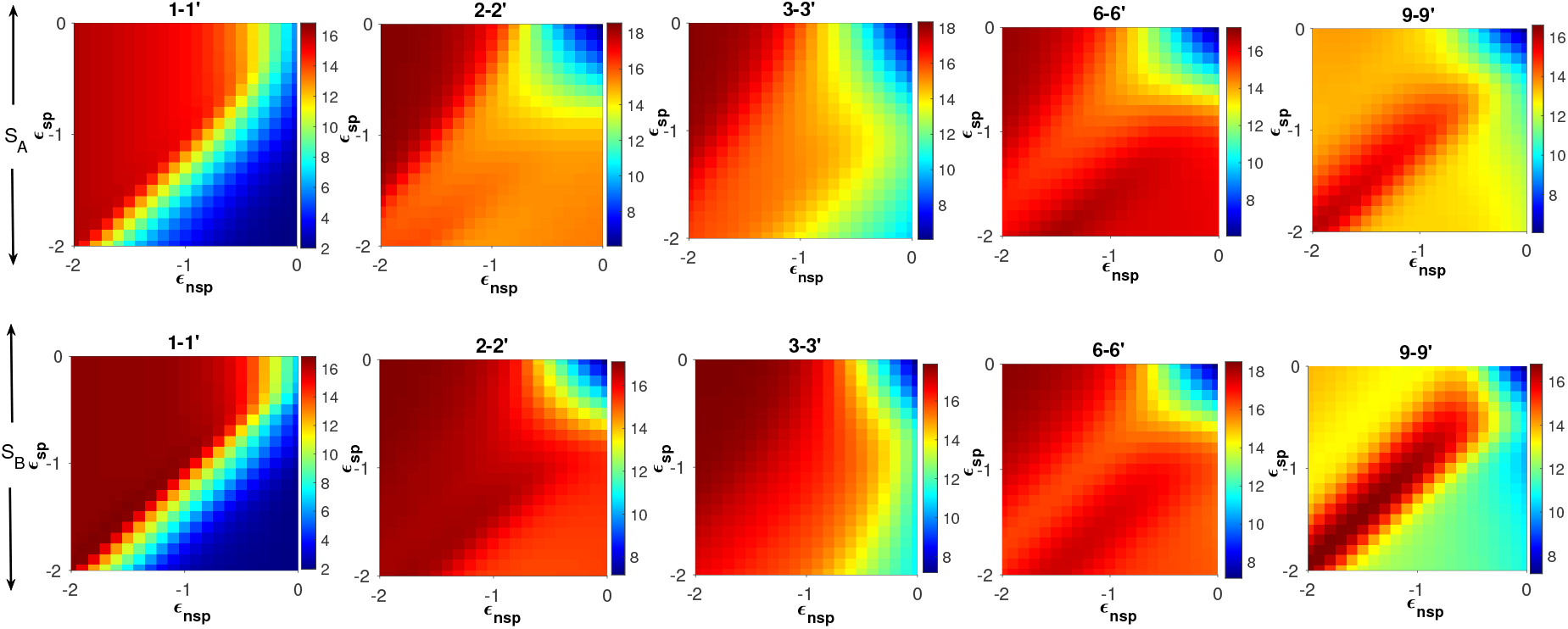
The horizontal upper panel shows total intra contacts (formed within polymer 1 and polymer 2) heatmap for shape *S*_*A*_ and lower pannel presents the total intra contacts heatmap as a function *ϵ*_*AA*_ and *ϵ*_*AB*_ for shape *S*_*B*_

As *ϵ*_*nsp*_ decreases and equals *ϵ*_*sp*_ = *ϵ*_*nsp*_ = −1, all polymer systems converge to a same separation time, behaving as homopolymers with very low segregation time, as they prefer to fold individually leading to less segregation time [21]. Further decreasing *ϵ*_*nsp*_ to −2, leads to increased segregation time for all the systems with maximum stable overlap in the 9-9’ system (see contact maps and heat map), resulting in the longest separation time for 9-9’ across both confinements. We also see that 1-1’ system start making individual non-specific contacts at higher non-specific strength which decreases the segregation time for 1-1’ at *ϵ*_*nsp*_ = −2.0. At further higher non-specific interaction strength *ϵ*_*nsp*_ = −3.0, 9-9’ system exhibits highest segregation time due to formation of lamellar structures, which are difficult to separate (see contact maps) [48, 49]. On the other hand, the 6-6’ system, which forms also lamellar like 𝒮-shaped structures and remains in close proximity throughout the non-specific interaction energy range, but encounters an entropic cost when trying to overlap two 𝒮-shaped polymers in lattice. This entropic barrier makes the 6-6’ system having moderate segregation time. The 2-2’ and 3-3’ systems fall between the extremes, neither fully overlapped nor entirely separated making frustrated phase locked configurations, with their block lengths and finite sizes individual folding/intra non-specific contact formation is easier for 3 and 3’ leading to lowest segregation time for 3-3’ system. As *ϵ*_*nsp*_ decreases, the segregation time of 1-1’ system decreases, with a noticeable reduction from a highly overlapped to a less overlapped state, though some finite contacts remain.

While the different polymer systems maintain the same order in segregation time across both square and rectangular confinements, the symmetric or square confinement (*S*_*A*_) leads to longer separation times compared to rectangular confinement (*S*_*B*_). This can be explained by the static study, which shows that the ground states in square confinement (*S*_*A*_) possess greater entropic freedom, allowing for more configurational possibilities. In contrast, the ground states in *S*_*B*_ are less degenerate and have higher free energy, limiting the configurations (as only linear configurations are preferred strongly). As a result, polymers take longer to separate within *S*_*A*_, likely due to higher entanglement in the more symmetric confinement. This High mixing of polymers in symmetric confinement is earlier seen in many studies [18, 21, 57]

The key takeaway from the separation time study is that, in the static analysis, among all the systems studied, the 9-9’ system is the hardest to separate at high non-specific interaction strength due to longer overlapped block size and all systems exhibit longer segregation time in symmetric confinement. As a limitation of the study, the 1-1’ system fails to accurately capture its dynamic behavior due to limitations of the lattice model. Simply put, as overlap between polymers increases, the free energy decreases, leading to longer segregation times.

## IV. CONCLUSION

In this paper, we investigated the static and dynamic aspects of segregation (or compaction) of two block co-polymers of varied block sizes subjected to varying non-specific (hetero-genetic) interactions under strong confinement, motivated by various studies of epigenetics driven chromosome organization by block copolymer modeling [7, 15, 40, 50]. The goal was to determine whether different block sizes of BCPs influence segregation or compaction depending on non-specific interactions, and whether the shape of the confinement accelerates or delays the segregation process. By employing a simple model, we also aimed to understand whether polymer systems exhibit behavior that is independent of the methods used, even for relatively short chain lengths like 18 where finite size effects are strong. As we summarize our study, we can confidently highlight two main findings. First, non-specific interaction energy plays a dominant role in the compaction process, particularly for polymers with larger block sizes such as 6-6’ and 9-9’. The 9-9’ polymers are especially difficult to separate due to the overlapping of different kind monomers forming lamellar phases [48, 49] (or “hugging” 𝒰 shapes). Lamella mediated compaction in chromatin organization is present in literature [54]. The phase separation induced self assembly compaction of long-range sequences driven by specific interaction strength directly corresponds to heterochromatin, where phase separated, inactive and stable domains are seen [15, 57, 58]. Additionally, short range sequences (2-2’ and 3-3’) experience frustration due to periodic stickers and spacers leading phase-locked meta stable states [44, 45]. Second, asymmetric or lateral confinement promotes greater segregation across the interaction range, as the polymers tend to form individual globules with more separation between them. Beyond considering the sequence information, the segregation time is lower within the lateral confinement *S*_*B*_ for all BCPs. This behavior bears a crude similarity to the polar organization of interphase sister chromatids, where they are pulled apart and move toward opposite ends of the deformed or laterally elongated cell [20]. As a limitation, the lattice representation of the 1-1’ system at low non-specific interaction strength fails to replicate its dynamic behavior, where it remains mostly separated. Where as the dynamic study reveals 1-1’ system due to its high entropic cost of overlapping remain decondensed or in coil shape corresponding to euchromatin which is typically active and less condensed [15, 57, 58].

DM would like to acknowledge the financial support provided by a Postdoctoral Fellowship at Lund University, Sweden, where part of this work was carried out. The support and resources provided by PARAM Shivay Facility under the National Supercomputing Mission, Government of India at IIT (BHU), Varanasi, are gratefully acknowledged. MD would like to acknowledge financial assistance from the SERB, India, UGC, India, SPARC scheme of MoE, and IoE scheme, MoE, India. DM and MD would like to thank Mamta Yadav for her valuable suggestions.

## APPENDIX: INDIVIDUAL CONTACT DISTRIBUTION

To enhance comprehensiveness, we plot the heat maps of total individual contacts (i.e., total contacts formed within polymer 1 and polymer 2) for all sequence cases within *S*_*A*_ and *S*_*B*_ in the upper and lower panels of 6. Mutual contacts which shows the compaction between two polymers where as individual contacts map shows the separation between the polymers. For 1 and 1’, the phase diagram exhibits a sharp transition at *ϵ*_*nsp*_ *< ϵ*_*sp*_ as the number of individual contacts *n*_*p*_ increases significantly. This behavior, described in the main text section, occurs because the individual contacts are mediated solely by *n*_*nsp*_ contacts.

The 2-2’ system maintains significant individual folding (≈13− 14) across the entire energy parameter range. Similar to the mutual contact map, the variation in individual contacts shows a more pronounced transition in *S*_*A*_ than in *S*_*B*_, possibly due to the symmetric confinement in *S*_*A*_, which allows for greater configurational entropy and more ground-state-like energy configurations. Additionally, at a specific *ϵ*_*sp*_, lowering *ϵ*_*nsp*_ causes 2 and 2’ to fold within *S*_*B*_ earlier than in *S*_*A*_. This suggests that lateral confinement may promote earlier folding.

In the case of 3 and 3’, for most of the energy phase diagram, the polymers are folded into itself except when *ϵ*_*nsp*_ ≈0. Similar to 2-2’ system, 3 and 3’ fold early within *S*_*B*_, potentially due to the lateral confinement shape aiding folding. Comparing with mutual contacts heat maps for the same 3-3’ system, it is evident that this system predominantly folds within itself, discouraging overlap.

Just as in mutual contact maps, individual contact maps also reveal that 6-6’ system is the most stable system in terms of total individual contacts (*n*_*p*_) variation as for the whole energy range *n*_*p*_ remains almost constant. So, 6-6’ system remains folded within themselves across a wide energy spectrum.

Lastly, the 9-9’ system exhibits a different behavior compared to all the other systems as described in main text. The 9 and 9’ maintain a moderate number of individual contacts throughout the energy range, except when *ϵ*_*sp*_ ≈*ϵ*_*nsp*_, where individual contacts jump to a maximum. In all systems, decreasing *ϵ*_*nsp*_ increases individual contacts, leading to a state where the polymer folds within itself, except in the 9-9’ system. In this system, individual folding predominantly occurs within the regime *ϵ*_*sp*_ ≈ *ϵ*_*nsp*_, and further decreasing *ϵ*_*nsp*_ reduces folding within itself as mutual overlapping increases (see 2). Thus, the 9-9’ system shows the highest degree of overlap at decreased *ϵ*_*nsp*_ while still maintaining finite individual contacts (*n*_*p*_ ≈ 12), indicating that 9 and 9’ may form a U shape, hugging each other.

